# Hydroxycitric acid inhibits oxalate nephropathies formation through crystallization regulation and activation of the PPARα pathway

**DOI:** 10.1101/2022.12.05.519215

**Authors:** Yi-Han Zhang, Shu-Jue Li, Bang-Xian Yu, Qing Liang, Xin-Yuan Sun

## Abstract

Oxalate-induced nephropathies comprise a range of kidney disorders, for which there are no efficient pharmacological treatments. Hydroxycitric acid (HCA) is a derivative of citric acid with a variety of pharmacological activities including reducing body weight and calcium salt deposition. However, the specific mechanism of inhibition of oxalate nephropathies by this compound is not well understood. In this study, we successfully applied bioinformatics-based and simulated drug molecular docking approaches to predict potential targets of HCA. Subsequently, we explored the molecular mechanisms of HCA inhibition of renal calcium oxalate (CaOx) deposition and nephrotoxicity in an oxalate-induced NRK-52E cell model and an oxalate nephropathy rat model. HCA could effectively inhibit CaOx crystal deposition and reduce crystal adhesion and oxidative damage, effectively inhibit lipid deposition caused by high oxalate, and reduce lipid nephrotoxicity. HCA is more effective than traditional stone medications in inhibiting CaOx deposition and kidney damage. Further cellular transcriptomic analysis and in vitro results showed that HCA could stably bind peroxisome proliferator-activated receptor α (PPARα) and promote PPARα-RXR heterodimer formation, thus promoting the expression of downstream oxidative stress molecules (Nrf2, HO-1, SOD) and inhibiting calcium ion release and mitochondrial dysfunction, thus reducing oxalate-induced renal lipid peroxidation damage. Therefore, HCA, a novel drug with the ability to modulate lipid metabolism and inhibit CaOx formation, may be a therapeutic option for the treatment of oxalate nephropathies.

## 1. Introduction

Kidney stones are one of the common urological diseases with a prevalence of 1.7-14.8% in different countries around the world [1]. The prevalence rate for Chinese adults is 5.8% (6.5% for men and 5.1% for women) [2]. High urinary oxalate levels are an important factor in oxidative damage to the kidney, and calcium oxalate (CaOx) is the most common crystalline component of kidney stones[3, 4]. Stone formation is inextricably linked to the body’s metabolic level, with risk factors including obesity, diabetes, hypertension, and metabolic syndrome [5]. Obesity has been shown to increase the risk of stone formation by 55% (95% CI: 1.25-1.94; P < 0.001) [1], so deranged lipid metabolism may be an important cause of stone formation. However, very little attention has been paid to the prevention of deranged lipid metabolism-induced stones.

Proximal tubular cells (PTC) are the cells with the highest energy requirements in the body [6]. Faced with this demand, these cells use fatty acids as their main source of energy, probably because fatty acids produce three times more adenosine triphosphate (ATP) than glucose [7]. Thus, dysregulation of lipid metabolism in PTC is thought to be closely correlated with dysfunction in PTC and ultimately leads to accelerated renal damage. A correlation between lipid nephrotoxicity and the progression of chronic kidney disease (CKD) has been suggested as early as 1982 [8]. Excessive accumulation of lipids in renal tubular cells can lead to lipotoxicity, causing lipid peroxidation damage to the cells. The pathogenic mechanism is an intracellular overload of free fatty acids, leading to an accumulated triglyceride pool, manifested by mitochondrial dysfunction, increased reactive oxygen species production and decreased ATP production, apoptosis, and elevated inflammatory cytokines [9, 10]. Oxidative damage to renal cells not only promotes CaOx formation, causing heterogeneous nucleation of crystals but also increases the adhesion of CaOx crystals to the cell surface [11].

Hydroxycitric acid (HCA) is a derivative of citric acid that is widely found in tropical rainforest plants such as Garcinia Cambogia and Hibiscus Subdariffa, and has been widely reported to have a weight and fat loss effect [12, 13]. HCA effectively inhibits the first step in fatty acid synthesis by inhibiting adenosine triphosphate citrate lyase and preventing the conversion of citric acid to acetyl coenzyme A [12]. HCA, therefore, plays a key role in inhibiting the synthesis of fatty acids, cholesterol, and triglycerides. HCA also has good antioxidant activity and can induce antioxidant gene expression by increasing Nrf2 nuclear expression and activating antioxidant response elements (ARE), thereby reducing oxidative damage in cells [13]. In addition, HCA can effectively complex calcium ions to inhibit the crystallization of CaOx. Chung et al. [14] showed that increasing the concentration of HCA inhibited CaOx monohydrate (COM) crystallization by 60%, reduced the rate of crystal growth as well as potentially inhibited COM nucleation and that HCA exhibited better crystallization inhibition than citric acid (CA). Meanwhile, Han and Liu et al. [15, 16] also conducted a preliminary study on the crystallization inhibition of K-HCA in mice and Drosophila high-oxalate models, and K-HCA could effectively inhibit the formation of CaOx crystals, but the molecular mechanism of stone formation inhibition by HCA has not been investigated in depth.

In this study, we investigated the role of HCA on the regulation of CaOx crystallization at the cellular and animal levels, and further obtained the potential targets of action and molecular mechanisms of HCA inhibition of oxalate nephropathies by RNA sequencing studies in hyperoxaluria molded rat kidneys. To further investigate the mechanism of action of HCA to exert pharmacological activity, the intermolecular interaction force between HCA and target genes was calculated by simulation to determine the binding ability of HCA to target genes and the site of action. Our study will provide further theoretical evidence for the clinical application of HCA as an inhibitory drug for oxalate nephropathies.

## 2. Materials and Methods

### 2.1 Materials

Human Renal Proximal Tubular Epithelial Cells (HK-2), Rat Renal Tubular Epithelial Cells (NRK-52E), Rat Normal Renal Fibroblasts Cells (NRK-49F), Madin-Darby Canine Kidney Cells (MDCK) were purchased from the Shanghai Cell Bank of the Chinese Academy of Sciences (Shanghai, China). The following materials were also purchased for the study: fetal bovine serum and cell culture medium (DMEM) (Gibco, USA), Oil Red O (sigma), Penicillin, and streptomycin (Solarbio, Beijing). The mitochondrial membrane potential assay kit with JC-1, rhodamine phalloidin, anti-fluorescence quenching sealer, Fluo-4/AM, Calcein AM/PI were purchased from the Shanghai Beyotime Bio-Tech Co., Ltd. (Shanghai, China). The cell proliferation assay kit (CCK-8) and reactive oxygen species (ROS) was purchased from KeyGEN Biotechnology Co. (Nanjing, China). Sodium oxalate (Na_2_Ox), potassium citrate (K_3_Cit-H2O), paraformaldehyde, and ethanol are analytically pure (Guangzhou Chemical Reagent Factory, China). Hydroxycitric acid (HCA) was purchased from Fortuneibo-Tech Co., Ltd (Shanghai, China). The triglyceride (TG) quantification kit was obtained from Elabscience Biotechnology Co., Ltd (Wuhan, China).

### 2.2 Experimental section

#### 2.2.1 Animal experiment

Sprague-Dawley (SD) male rats, about 6-8 weeks old, weighing about 180-220 g, were purchased from Guangdong Medical Laboratory Animal Center. 32 SD male rats were randomly divided into four groups as follows: (A) normal control group (Control), normal chow feeding; (B) stone group (EG): fed normal chow, drinking 1% ethylene glycol by volume for 28 days, and treated accordingly with 1% ammonium chloride by gavage for the first three days of the first and third weeks of the experiment; (C) HCA-treated group (HCA), treated with 0.1 mmol/kg body weight of HCA by gavage daily for 28 days in parallel with the corresponding treatment in the stone group; (D) K_3_Cit-treated control group (K_3_Cit), treated with 0.1 mmol/kg body weight of K_3_Cit by gavage daily for 28 days in parallel with the corresponding treatment in the stone group. All rats were allowed to eat regular food ad libitum and were kept at 25°C and under a light/dark cycle for the duration of the experiment. All procedures were carried out following the animal management regulations of the Ministry of Health of the People’s Republic of China and approved by the Animal Care Committee of the First Affiliated Hospital of Guangzhou Medical University.

#### 2.2.2 Observation of renal HE staining and CaOx crystal deposition

Rat kidneys were isolated and sectioned, fixed in 4% paraformaldehyde, and then embedded in paraffin. Cross-sectional sections of the kidneys (4 μm) were prepared and stained with hematoxylin-eosin. We then scored pathologically the tubular injury in rats by reference to the method of Li et al. [17]. The tubular injury was assessed by luminal dilatation, epithelial cell necrosis and loss of brush border. Five fields of view were randomly selected for scoring under a 400× microscope and graded according to the area of injury (grade 0, no injury; grade 1, <20% injury; grade 2, 20-40% injury; grade 3, >40% injury), and the histopathological changes were scored semi-quantitatively in a blinded manner. Sections were then analyzed for crystal deposition by polarized light optical microphotography (CX31-P, Olympus, Japan). The size of the crystal deposition area was determined by ImageJ software to quantify the number and area of CaOx crystal deposits in the renal tissue sections.

#### 2.2.3 Periodic Acid-Schiff stain (PAS) staining

Renal tissue sections were stained with PAS to observe the integrity of the glomerular basement membrane and thylakoid matrix and to evaluate the tubular injury fraction according to the literature [18]. At least three sections were used in each group, and five randomly selected fields of view (×200 magnification) were used for each section to calculate the tubular injury fraction.

#### 2.2.4 Immunohistochemical staining

Immunohistochemical staining of kidney sections was performed according to the literature [19]. Kidney sections were incubated overnight at 4°C using antibodies against Anti-FABP4 (1:2000 for IHC, Proteintech), anti-ACOX-1 (1:250 for IHC, Proteintech), anti-OPN (1:500 for IHC, Proteintech), and anti-IL-6 (1:500 for IHC, Proteintech), Anti-PPARα (1:100 for IHC, Affinity), Anti-Nrf-2 (1:200 for IHC, Abclonal), Anti CD44 (1:500 for IHC, Abcam). Images were acquired using a PathScope™ 4S scanner (DigiPath, USA) and quantified using Image J software.

#### 2.2.5 RNA sequencing and enrichment analyses

Three parallel groups were set up for each of the normal control groups (Control), stone groups (EG), and HCA treated groups (HCA). Total RNA was isolated from the Control, EG, and HCA groups using Trizol RNA reagent (Invitrogen) according to the manufacturer’s instructions. cDNA libraries were constructed and transcriptome sequencing (RNA seq) was performed using the Illumina HiSeqTM 2500 provided by Gene Denovo Biotechnology Co. (Guangzhou, China). To determine differentially expressed genes (DEGs), the “DESeq2” package by R soft was used based on p < 0.05. Heat maps were generated using R soft. Kyoto Encyclopedia of Genes and Genomes (KEGG) pathway analyses of DEGs were conducted using the “cluster profile” package.

#### 2.2.6 Cell culture and transfection

Cell culture: NRK-52E cells were cultured in DMEM medium containing 10% fetal bovine serum, penicillin-streptomycin antibiotics, and placed in a constant temperature incubator at 37°C and 5% CO_2_.

Cell transfection: the small interfering RNA (siRNA) was synthesized by Tsingke Biotechnology Co. The following siRNA sequences were used in the study: R25747-siPPARa-1 (On-Target Plus: 5’-CGAUCAAUAAUUUCAGACU-3’ and 5’-UAUUCGACACACGAUGUUCTT-3’), R25747-siPPARa-2 (On-Target Plus: 5’-GGCGAACUAUUCGGCUAAATT-3’ and 5’-UUUAGCCGAAUAGUUCGCCTT-3’), and R25747-siPPARa-3 (On-Target Plus:5’-CGAUCUGAAAGAUUCGGAATT-3’ and5’ - UUCCGAAUCUUUCAGAUCGTT-3’). After transfection with Lipofectamine 3000 and the indicated plasmids or siRNAs for 48 h according to the manufacturer’s instructions (Invitrogen), the transfection efficiency was assessed by Western blotting (WB) to detect protein levels. The candidate siRNA with the highest knockdown efficiency was then selected for subsequent experiments.

#### 2.2.7 Assay of HCA on the viability of different types of kidney cells

Cell suspensions (1 × 10^5^cells/mL) were inoculated in 96-well plates at 100 μL per well and incubated for 24 h. Serum-free medium containing HCA at concentrations of 0.125, 0.25, 0.5, 0.75, 1, 1.25, 1.5, 1.75, and 2 mmol/L was co-cultured with four types of renal cells (HK-2, NRK-52E, MDCK, NRK-49F) for 24 hours. Subsequently, 10 μL of CCK-8 reagent was added to each well and placed in the incubator for 2 h. The change in cell viability was calculated by measuring the OD at 450 nm with an enzyme marker.

#### 2.2.8 Cytotoxicity assay of oxalate

Cell suspensions (1 × 10^5^cells/mL) were inoculated in 96-well plates at 100 μL per well and incubated for 24 h. Serum-free medium containing oxalate at concentrations of 0.25, 0.5, 0.75, 1, 1.25, 1.5, 1.75, and 2 mmol/L was co-cultured with NRK-52E cells for 24 h, and then OD values were measured to calculate cell viability.

#### 2.2.9 Detection of the cell viability of HCA treated oxalate-damaged cells

Cell suspensions (1 × 10^5^cells/mL) were inoculated in 96-well plates with 100 μL per well and incubated for 24 h. 1 mmol/L oxalate solution was selected for damaged NRK-52E cells to construct the damage cell model, and the concentrations of 1, 2, and 4 mmol/L HCA solution were added and co-incubated for 24 h. The OD values were determined according to the instructions of the CCK-8 kit, and the Cell viability was calculated.

#### 2.2.10 Observation of CaOx crystallization regulation by HCA

The cells were incubated for 0.5 h, 2 h, and 24 h in a 12-well culture plate, and serum-free DMEM medium containing 1 mmol/L oxalate was added to each well, and the cells were treated with HCA at concentrations of 0.5, 1 and 2 mmol/L, respectively. The size, shape, and distribution of the formed crystals and the state of the cells were observed by an ordinary microscope.

#### 2.2.11 Cytoskeleton observation

1 mL of cell suspension (1 × 10^5^cells/mL) was inoculated in 12-well culture plates for 24 h. The experiments were divided into the following four groups: (1) normal control group, which was added serum-free medium only and incubated for 24 h; (2) HCA control group, which was treated with serum-free medium containing 2 mmol/L HCA for 24 h; (3) oxalate damaged group, serum-free medium containing 1 mmol/L oxalate was added and incubated for 24 h; (4) HCA-treated group, serum-free medium containing 1 mmol/L oxalate and 2 mmol/L HCA solution was added and incubated for 24 h. After reaching the treatment time, the cell culture solution was aspirated, and fixed by 4% paraformaldehyde at room temperature for 20 min. The cells were then permeabilized with 0.1% Triton X-100 for 5 min, stained with rhodamine-phalloidin, placed in an incubator at 37°C for 1 h. The whole process was operated avoiding light, and the nuclei were stained with DAPI for 5 min and observed under a fluorescence microscope.

#### 2.2.12 Detection of intracellular reactive oxygen species (ROS) level

The experiments were grouped as in 2.2.11. After reaching the treatment time, 500 μL of DCFH-DA staining solution diluted with the serum-free medium was added to each well for 30 min, washed 3 times with PBS, placed under a fluorescent microscope protected from light to observe the intracellular ROS levels, and the fluorescence intensity was counted by ImageJ software.

#### 2.2.13 Intracellular calcium level assay

After reaching the treatment time, the culture solution was aspirated, the cells were washed 3 times with PBS, 2 μmmol of Fluo-4 AM assay working solution diluted with 500 μL of culture medium was added, incubated for 30 min at 37°C, washed 3 times with PBS, placed under a fluorescent microscope to observe the intracellular calcium ion distribution, and the fluorescence intensity was counted by ImageJ software.

#### 2.2.14 Detection of mitochondrial membrane potential (ΔΨm)

After reaching the treatment time, the culture solution was aspirated, the cells were washed 3 times with PBS, 200 μL of JC-1 staining solution was added for 60 min, the cells were washed 3 times with PBS, and the ΔΨm was observed under a fluorescence microscope, and the fluorescence intensity was counted by ImageJ software.

#### 2.2.15 Live/dead cell double staining assay

After reaching the treatment time, the culture medium was aspirated, the cells were washed twice with PBS, the appropriate volume of Calcein AM/PI assay working solution was added, incubated for 30 min at 37ºC and protected from light, the cells were washed three times with PBS and placed under a fluorescence microscope to observe the fluorescence staining (green fluorescence for Calcein AM, Ex/Em=494/517 nm; red fluorescence for PI, Ex/Em=535/617 nm). The green fluorescence markers are live cells stained with Calcein, and the red fluorescence markers are dead cells stained with PI.

#### 2.2.16 Lipid deposition staining observation

Oil red O staining was used to observe the cellular lipid deposition. 2 mL of cell suspension with a cell concentration of 1.0×10^5^ cells/mL was inoculated in a 6-well culture plate containing cell slides for 24 h. After reaching the treatment time, the slides were fixed in 10% formalin for 10 min and rinsed 3 times in PBS (pH=7.4). The slides were then placed in 1% Oil Red O staining solution dissolved in 60% isopropyl alcohol and left at room temperature for 20 min. 70% ethanol was used to wash the slides for 2 s to remove background staining and hematoxylin staining was performed for 10 s. The samples were placed under a general microscope to observe lipid deposition and quantify it with ImageJ software.

#### 2.2.17 Western blot (WB) experiments

To obtain total protein, NRK-52E cells were treated with RIPA lysis buffer. Protein samples were separated on SDS-PAGE and proteins were transferred to polyvinylidene difluoride (PVDF) membranes. PVDF membranes were incubated in a blocking buffer containing 5% (w/v) bovine serum albumin (BSA). Protein extract was incubated overnight at 4°C with primary antibodies against PPARα (1:1000 for WB, Proteintech), anti-FABP4 (1:1000 for WB, Proteintech), anti-αRXR (1:1000 for WB, Proteintech), anti-SOD (1:1000 for WB, Proteintech), anti-HO -1 (1:1000 for WB, Proteintech), and GAPDH (1:1000 for WB, Proteintech), and Nrf2 (1:1000 for WB, Abclonal). The PVDF membranes were then incubated with HRP-coupled secondary antibodies (1:10,000, CST) for 2 h at 25°C and visualized by ChemiDoc XRS (Bio-Rad) instrument.

#### 2.2.18 Component-target molecule docking

The 2D structure of HCA with fenofibrate was obtained from the PubChem database (https://pubchem.ncbi.nlm.nih.gov/), converted to mol2 format by Chem3D, and then saved as a docking ligand in PDBQT format. The UniProt database (https://www.uniprot.org/) was searched for the target protein with good resolution: PPARα. To perform molecular docking, the ligands need to be optimized for energy minimization and the pdbqt format file needs to be constructed using the Autodock docking tool [20]. Autodock can be used to simulate the docking of a drug to a target protein. A docking fraction less than 0 means that the ligand can spontaneously bind to the receptor [21].

#### 2.2.19 Triglyceride (TG) measurement

After reaching the treatment time, the treated cells were collected for ultrasonication to sufficiently break the cells. The cells were centrifuged at 10,000g for 10 min at 4°C. The enzyme working solution was added to generate red quinones of the benzoquinone imide family, the color shade of which was proportional to the TG content, and the OD value of the supernatant at 510 nm was measured by an enzyme marker.

#### 2.2.20 Statistical analysis

Experimental data were expressed as mean ± standard deviation (x ± SD). The experimental results were statistically analyzed using Graphpad Prism 9.0 software, and mean differences were compared using ANOVA and Student’s t-test, and a p-value less than 0.05 was considered statistically significant.

## 3. Results

### 3.1 HCA inhibits oxalate-induced CaOx deposition and cellular injury in rat kidney

To investigate the inhibitory effect of HCA on oxalate-induced CaOx deposition and cell damage, we verified the effect of HCA *in vivo* in an ethylene glycol-induced high oxalate model in rats. The HE staining results showed that the kidneys of the normal control rats were of good size and shape, with regular glomerular morphology and tightly arranged tubules. The kidneys of the stone group showed structural damage such as dilatation, destruction, discontinuity, or even breakage of the tubules and loss of epithelial cells (Figure 1A). Pathological scoring of renal tubular injury in rats was performed by the results of HE staining (Figure 1D). Compared to the normal group, the stone group had a significantly higher tubular injury score (p<0.001) and a severe degree of injury and inflammatory infiltration. After HCA intervention, tubular destruction in the kidney was significantly reduced (p<0.01) and significantly better than in the potassium citrate-treated group (K_3_Cit, p<0.05). PAS staining was further used to assess tubular injury of renal tissue sections (Figures 1B&1E). The stone group had severe tubular destruction and significantly higher scores, while the HCA-treated group had normalized renal morphology and intact renal units. In contrast, PAS scores in the K_3_Cit group were not significantly different from those in the modeling group.

**Figure 1.**
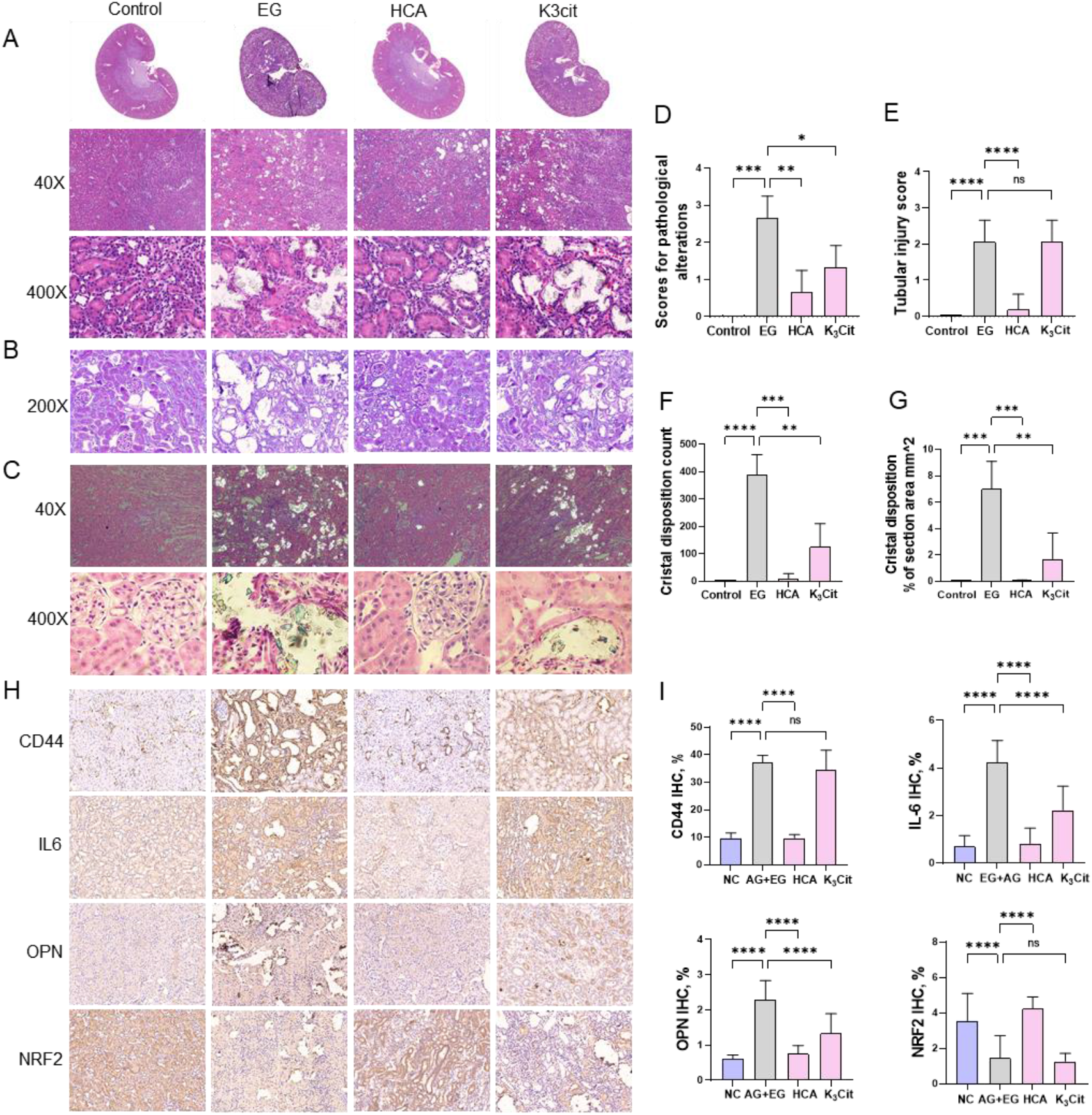
HCA inhibits CaOx deposition and cell injury in rat kidneys induced by oxalate. (A) HE staining of the kidney; (B) PAS staining; (C) polarized light microscopy; (D) renal tubular injury score based on HE staining; (E) injury score based on PAS; (F) crystal number statistics; (G) crystal area statistics; (H) immunohistochemical staining results of CD44, OPN, IL-6 and Nrf2 proteins in kidney tissues, with the magnification of 200X, Scale bar=50 μm; (I) semi-quantitative results of CD44, OPN, IL-6, Nrf2 expression levels by ImageJ for mean optical density semi-quantification. * indicates P < 0.05, ** indicates P < 0.01, *** indicates P < 0.001, and **** indicates P < 0.0001.

HCA significantly inhibited the formation of CaOx crystals, in addition to inhibiting oxalate-induced kidney injury. Based on the birefringent nature of CaOx crystals, we used polarized light microscopy to observe the formation of CaOx crystals in rat kidneys (Figure 1C). The CaOx crystals in the stone group were distributed in the tubular lumen of the renal cortex and the renal medulla. A large number of rhombic CaOx crystals of different sizes and sharp edges adhered to the surface of the tubular epithelium in clusters. Some of the crystals are embedded in the tubular lumen, causing dilatation, destruction, and discontinuity of the tubules or even rupture. The destruction of the tubular lumen and glomeruli in the kidneys of the HCA-treated group was slight, with only a small amount of CaOx crystal formation and a 97.3% decrease in crystal deposition compared to the stone group (Figure 1F), which was significantly better than the K_3_Cit group (p<0.01).

We analyzed the expression of injury and adhesion-related proteins in the kidney by immunohistochemistry (Figure 1H). The expression of the antioxidant molecule-Nrf2 protein was significantly decreased (p<0.0001) and the expression of the adhesion molecules CD44, OPN, and the inflammatory molecule IL6 was significantly increased (p<0.0001) in the stone group of rats compared to the normal control group. HCA significantly reversed the alterations in the expression of these proteins and was obviously more effective than the K_3_Cit treatment group. These results suggest that HCA treatment significantly attenuates CaOx crystal-induced oxidative stress damage and inflammation and adhesion risk *in vivo*.

### 3.2 HCA prevented the changes of oxalate-induced cellular transcriptomic *in vivo*

To unravel the molecular mechanism by which HCA inhibits oxalate nephropathy, we performed RNA sequencing (RNA-seq) analysis. A total of 569 differentially expressed genes (DEGs) were identified between the stone group, the normal control group, and the HCA-treated group (Figure 2A). RNA-seq of renal tissue showed a similar gene expression profile between the normal control group and the HCA-treated group, while the stone group induced drastic changes. The result indicates that oxalate-induced cellular transcriptomic changes are prevented by HCA *in vivo*. Gene set enrichment analysis of differently expressed genes in the stone group versus the HCA-treated group revealed the altered gene expression mainly involved in lipid metabolism and oxidative stress responses, such as the peroxisome proliferator-activated receptor (PPAR) signaling pathway, adipocytokine signaling pathway, fatty acid degradation, and peroxisome pathway (Figures 2B&2C). The transcript levels of PPAR signaling pathway-related molecules were significantly higher in the HCA-treated group than in the stone group (Figure 2D), and the PPARα gene expression was most significantly altered. We speculate that HCA may play a role in protecting the kidney by activating the PPARα signaling pathway. In addition, the overall trends of differential genes in lipid metabolism-related pathways, inflammation and adhesion-related pathways, and oxidative stress-related pathways were found to be consistent in the GSEA analysis of the stone group versus the HCA-treated group (Figure 2E), suggesting that oxidative stress induced by high oxalate may be related to lipid metabolism pathways.

**Figure 2.**
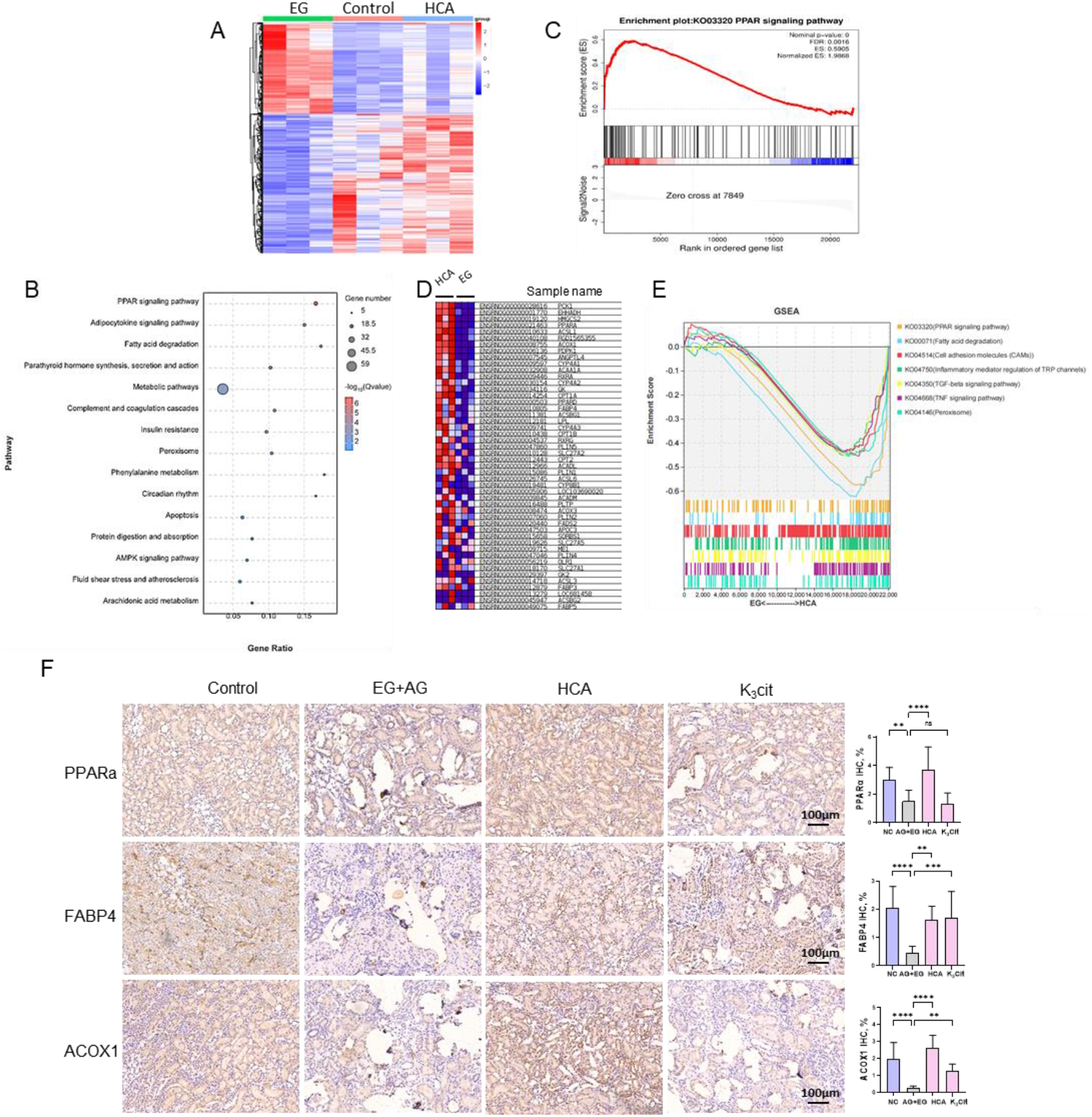
HCA prevented oxalate-induced cellular transcriptomic changes *in vivo*. (A) Heatmap of relative gene expression levels determined by RNA sequencing. Samples were analyzed in triplicate. (**B**) GSEA evaluated the enrichment of the PPAR signaling pathway in the HCA-treated group compared with the EG group. (**C**) Results of KEGG analysis of DEGs between EG and HCA treated group showed the biological processes. (**D**) The levels of expression of PPAR-associated genes were shown as a heat map; upregulated and downregulated genes are shown in red and blue, respectively. (**E**) GSEA was performed to evaluate the enrichment of gene set-associated metabolic pathways and cell damage pathways in EG and HCA-treated groups. (F) Semi-quantitative statistical plots of immunohistochemical staining results of PPARα, FABP4, and ACOX1 proteins and related expression levels at 100X magnification, Scale bar=100 μm. ** indicates P<0.01, *** indicates P<0.001, **** indicates P<0.0001.

To verify whether HCA protects the kidney by activating the PPARα signaling pathway, we examined the expression levels of PPARα, FABP4, and ACOX1 proteins in rat kidneys by immunohistochemistry (Figure 2F). The IHC assay showed that the expression of PPARα, FABP4, and ACOX1 was significantly decreased in the kidneys of the stone group compared to the normal group, while the expression was increased in the HCA-treated group. The results suggest that HCA may reduce oxalate-induced lipotoxic injury by activating the PPARα pathway.

### 3.3 Effect of HCA on cell viability and CaOx crystallization

To assess the safety of HCA, we assayed the cytotoxicity of HCA at concentrations ranging from 0.125 to 2.0 mmol/L in different types of renal cells, including NRK-52E, HK-2, MDCK, and NRK-49F cells, using the CCK-8 kit. At concentrations below 2 mmol/L, HCA had little effect on the viability of the four renal cells, indicating that HCA is safe below 2 mmol/L concentrations (Figures 3A-3D).

**Figure 3.**
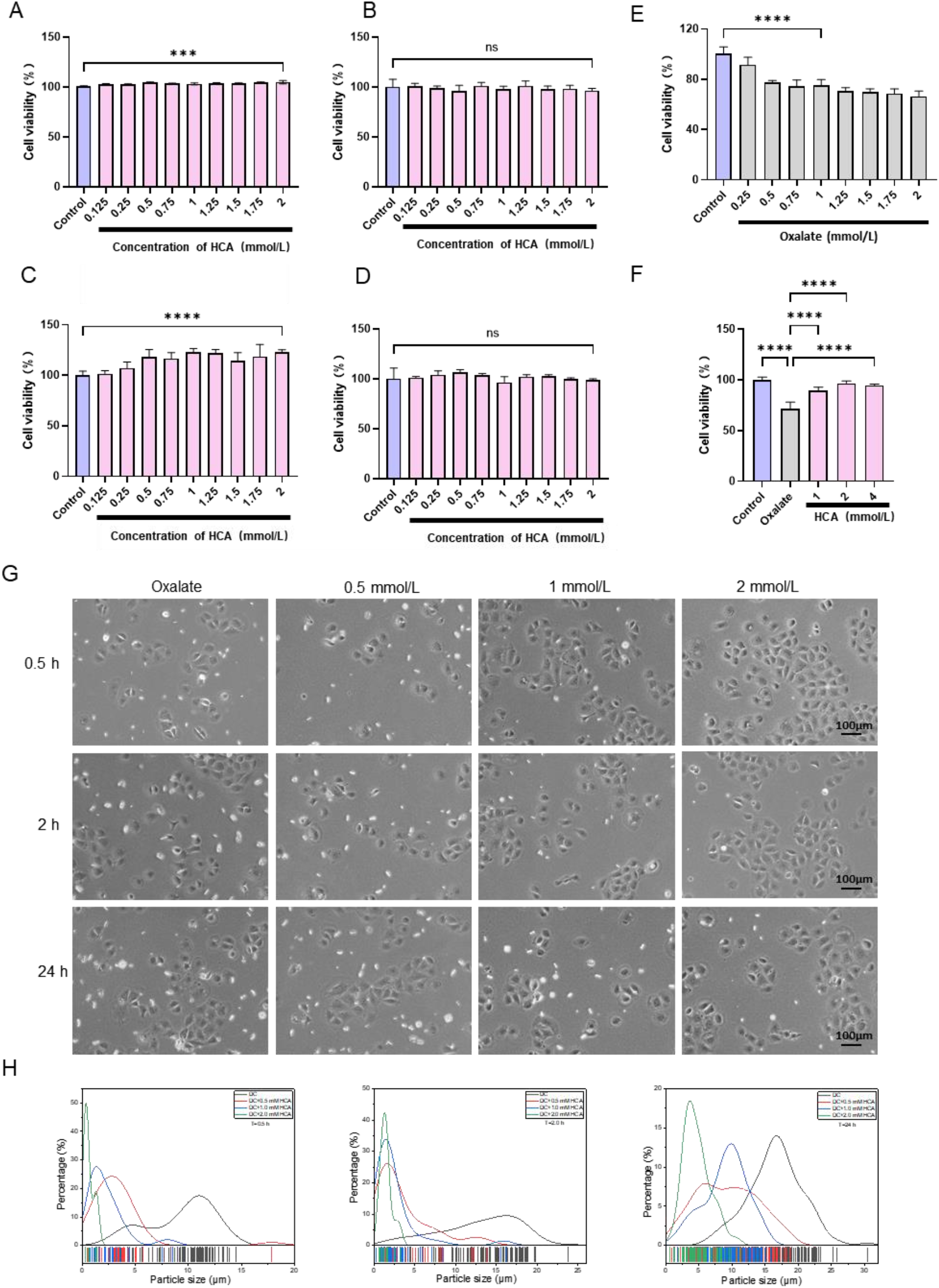
Effect of HCA on cell viability and calcium oxalate crystallization. (A-D) Effects of different concentrations of HCA on the activities of HK-2, NRK-52E, MDCK, and NRK-49F cells. HCA concentration: 0.125-2mmol/L; (E) effects of different concentrations of oxalate on the activities of NRK-52E cells, oxalate concentration: 0.25-2.0 mmol/L; (F) effects of different concentrations of HCA on oxalate damaged NRK-52E cells. Oxalate concentration: 1.0 mmol/L, HCA concentrations: 1, 2, and 4 mmol/L. (G) Effect of HCA on oxalate-induced CaOx crystallization; Oxalate concentration: 1 mmol/L, HCA concentration: 2 mmol/L; Crystallization time: 0.5 h, 2 h, 24 h. (H) The statistical plot of the variation of the particle size of CaOx crystals formed at different time points and different HCA concentrations. Scale bar = 100 μm. *** indicates P < 0.001; **** indicates P < 0.0001; ns indicates P > 0.05, no statistical difference.

Meanwhile, we examined the effect of different concentrations of oxalate on the viability of NRK-52E cells. The cell viability decreased gradually with increasing oxalate concentration and was 75.25% at a concentration of 1.0 mmol/L (Figure 3E). We also examined the effect of HCA at concentrations of 1.0-4.0 mmol/L on the change in cell viability caused by 1.0 mmol/L oxalate, and HCA at 2.0 mmol/L significantly inhibited the decrease in cell viability caused by oxalate (Figure 3F).

We observed the growth of CaOx by ordinary microscopy at different concentrations of HCA and different crystallization times (Figure. 3G). After the addition of 1 mmol/L oxalate, more CaOx particles were formed at 0.5 h and were present as a large amount of hexagonal rhombic CaOx monohydrate (COM) and a small amount of tetragonal bipyramid CaOx dihydrate (COD). As the crystallization time increases, the number of crystals formed gradually becomes greater and their size gradually becomes larger. At the same time, the state of the cells gradually deteriorates and becomes disordered as more crystals are formed on the cell surface. With the intervention of 0.5, 1, and 2 mmol/L HCA, the nucleation time of CaOx became progressively longer and the number of crystals formed became progressively smaller (Figure. 3H). In particular, at the 2 mmol/L concentration, only a few fine crystals were formed even at 24 h. The CaOx formed was mainly in the form of COD and the cell state was significantly improved. This suggests that HCA can significantly prolong the nucleation time of CaOx crystals and inhibit the growth of CaOx crystals, thus reducing the damage of NRK-52E cells by high oxalate.

### 3.4 HCA significantly inhibits oxalate-induced cell damage

We fluorescently stained the cells for F-actin by FITC-phalloidine and observed changes in cytoskeleton and cell morphology (Figure 4A). Actin is an important protein fibril in the cytoskeleton and plays an important role in maintaining the morphological stability of the cell. In the control group, the actin microfilaments were well-structured and the cell morphology was spacious and full. In the oxalate-damaged group, the microfilaments were disorganized, the cells were crumpled, the microfilament skeleton was blurred and the nuclei were brightened. In the HCA-treated group, it was observed that the microfilaments of the cells gradually regained their clarity and the cell morphology became more full-bodied.

**Figure 4.**
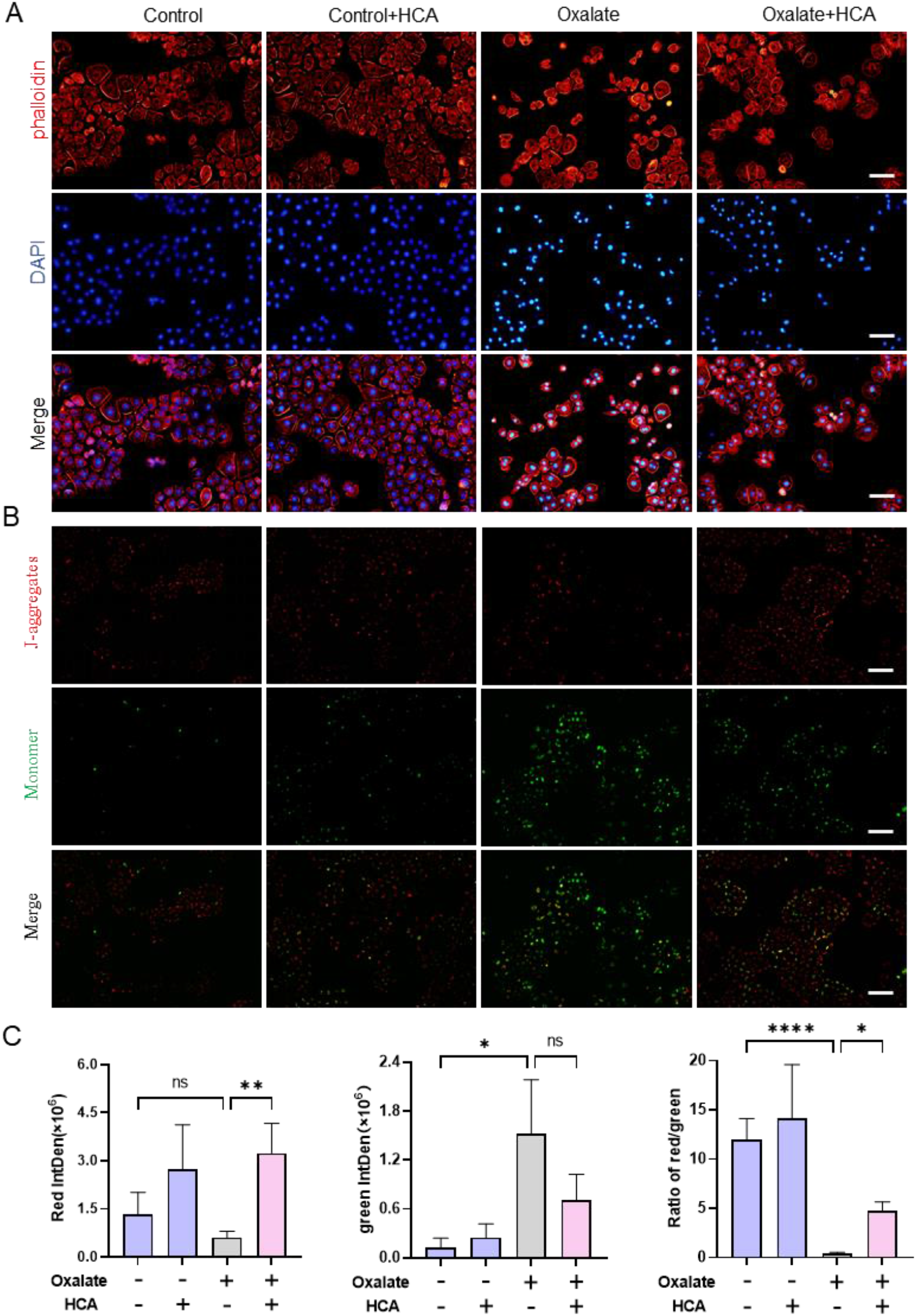
HCA inhibits the changes in cytoskeleton and mitochondrial membrane potential induced by high oxalate. (A) Fluorescence microscopy of the cytoskeleton and cell morphology changes of NRK-52E induced by oxalate. The nucleus (blue), and cytoskeleton (F-actin, red) were stained by DAPI and FITC-phalloidine, respectively. Magnification is 200X, Scale bar=100 μm. (B) Fluorescence microscopy of mitochondrial membrane potential with green fluorescence of monomer of JC-1; J-aggregates as aggregates of JC-1 in the mitochondrial matrix, red fluorescence; Merge as a combined plot of green fluorescence and red fluorescence; Magnification is 100X, Scale bar=200 μm. (C) semi-quantitative histogram of red fluorescence, green fluorescence, and red fluorescence to green fluorescence ratio of JC-1; oxalate concentration: 1.0 mmol/L; HCA concentration:2.0 mmol/L; treatment time: 24 h. * indicates P < 0.05, ** indicates P < 0.01, **** indicates P < 0.0001, ns indicates P > 0.05.

We analyzed the effect of HCA on mitochondrial membrane potential in oxalate-damaged NRK-52E cells by fluorescent probe JC-1 labeling staining (Figures 4B&4C). It shows strong red fluorescence and weak green fluorescence in normal NRK-52 cells. Upon exposure of the cells to the oxalate solution, the mitochondrial membrane underwent depolarization, resulting in a decrease in red fluorescence intensity and an increase in green fluorescence intensity. The red fluorescence intensity became stronger and the green fluorescence intensity was significantly reduced under the treatment of HCA, which improved mitochondrial membrane depolarization in epithelial cells and restored mitochondrial function.

Excessive damage to the cell disrupts the integrity of the mitochondrial membrane, causing the release of Ca^2+^ from the mitochondria, resulting in an increase in cytoplasmic Ca^2+^ content, which in turn causes oxidative stress in the cell. We labeled intracellular Ca^2+^ by Fluo-4/AM fluorescent probes and observed changes in fluorescence intensity by fluorescence microscopy (Figure 5A). In the normal group, the green fluorescence intensity was weak and only a small amount of Ca^2+^ was observed in the cytoplasm. In the oxalate-damaged group, the intra- and extracellular Ca^2+^ homeostasis was disrupted, the green fluorescence intensity was enhanced and the intracellular Ca^2+^ content was significantly increased. The green fluorescence intensity of the HCA-treated group was reduced and close to that of the normal group, indicating a decrease in intracellular Ca^2+^ levels, suggesting that HCA inhibited the oxalate-induced increase in intracellular Ca^2+^ concentration (p<0.0001, Figure 5D).

**Figure 5.**
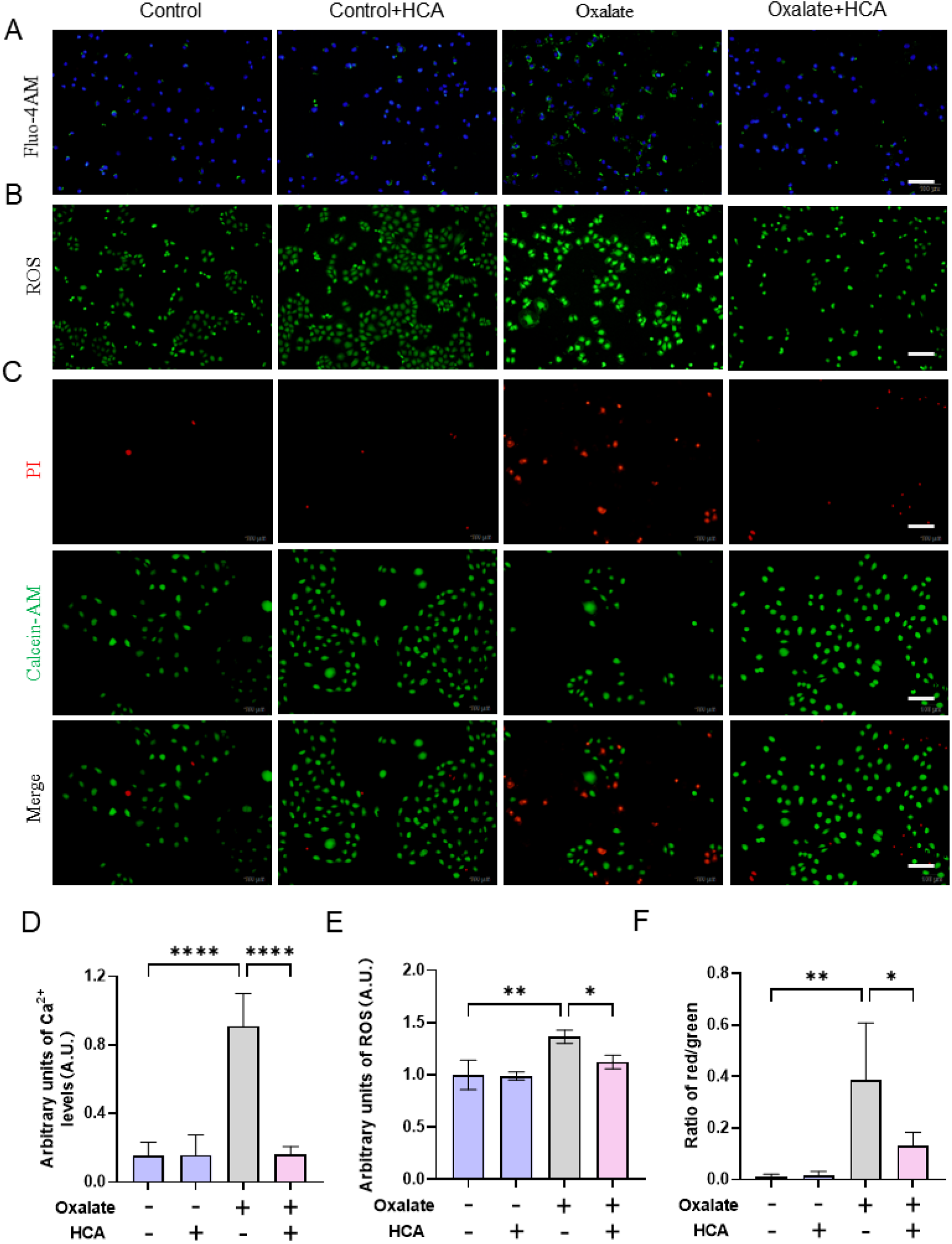
HCA inhibits cell damage induced by oxalate. (A) Fluorescence microscopy observation of intracellular Ca^2+^ level changes; (B) fluorescence microscopy observation of reactive oxygen species level changes; (C) fluorescence microscopy observation of live/dead cell staining with Calcein AM staining live cells with green fluorescence and propidium iodide (PI) staining dead cells with red fluorescence; (D-F) statistical results of intracellular Ca^2+^ level, ROS level and live/dead cell Statistical results of intracellular Ca^2+^ level, ROS level and the number of living dead cells. Oxalate concentration: 1 mmol/L; treatment time: 24 h; HCA concentration: 2 mmol/L. Magnification is 200X, Scale bar = 100 μm; * indicates P < 0.05, ** indicates P < 0.01, **** indicates P < 0.0001, ns indicates P > 0.05, no statistical difference.

In addition, high oxalate also caused intracellular oxidative stress damage, resulting in elevated intracellular ROS levels (Figures 5B&5E). The ROS levels were significantly reduced in HCA-intervened cells (p<0.05). Excessive ROS production can cause cell death. We observed the number of live and dead cells by Calcein AM/PI fluorescent double staining (Figure 5C). The normal group of NRK-52E cells had a good outline morphology with strong green fluorescence and only very few dead cells. After the oxalate treatment, the number of cells was significantly reduced and the proportion of dead cells was significantly higher compared to the normal group (p<0.01). After HCA intervention, the ratio of dead cells/live cells was reduced by approximately 66.5% compared to the oxalate group (p<0.05, Figure 5D), indicating that HCA was able to attenuate the oxidative damage to NRK-52E cells caused by oxalate and reduce cell death.

### 3.5 High oxalate induces lipotoxicity and damage in NRK-52E cells

We used high oxalate to damage NRK-52E cells to construct a model of damaged cells and observed intracellular lipid accumulation by oil red staining (Figure 6A). The results showed that normal cells had fewer intracellular lipids. As oxalate concentration increased, intracellular lipid accumulation gradually increased, cells were wrinkled and deformed, and cell damage increased (Figure 6D). In addition, oxalate reduced the expression of lipid metabolism-related molecules - PPARα, fatty acid binding protein 4 (FABP4), retinoid X receptor (RXR), and antioxidant molecule - Nrf2 in a dose-dependent manner (p<0.05) (Figures 6B&6C). High oxalate causes a disturbance in lipid metabolism and further decrease cellular antioxidant ability. Overload of intracellular free fatty acid accumulation leads to the accumulation of intracellular triglycerides and increased production of ROS, which can cause damage to cells [10].

**Figure 6.**
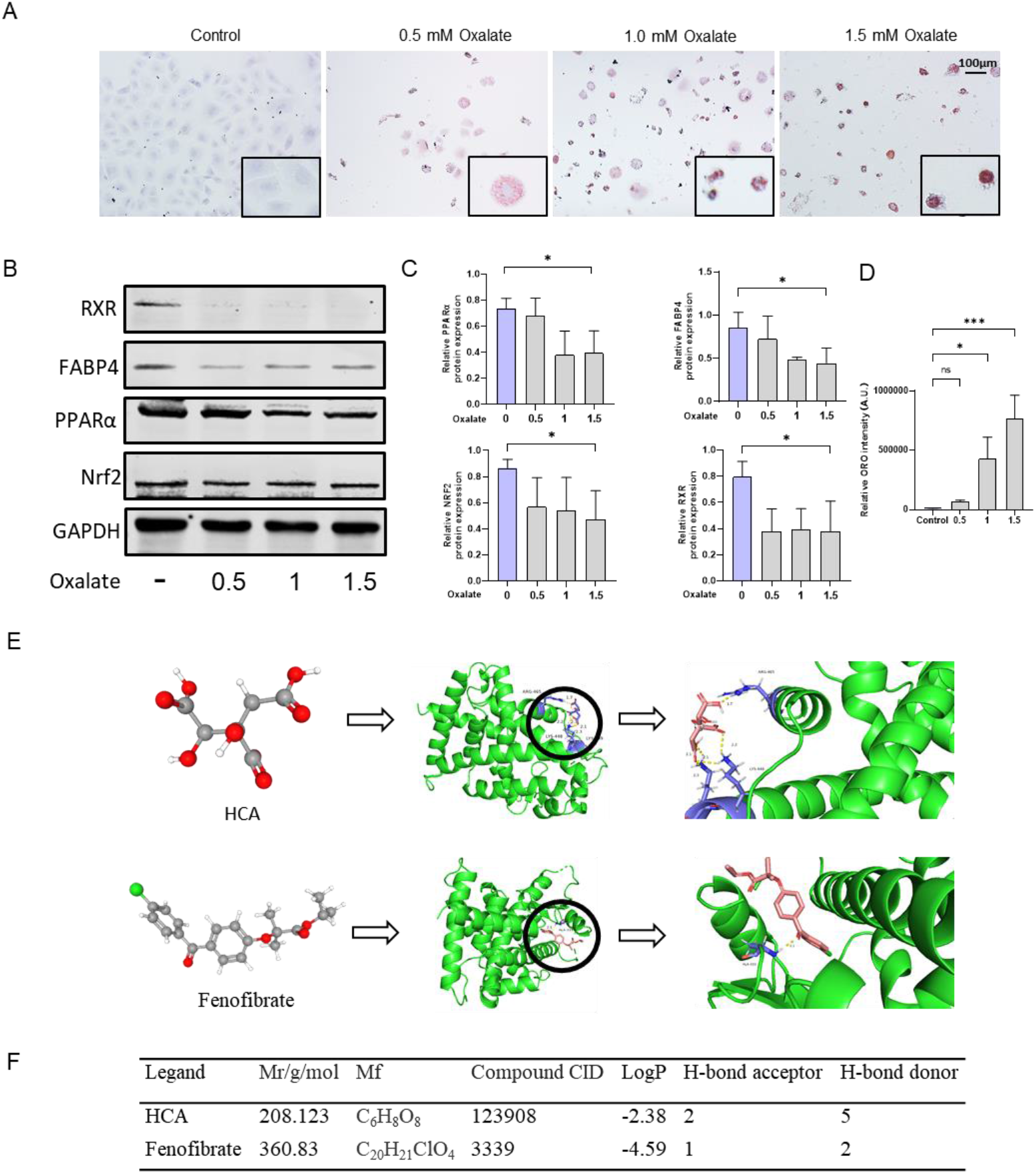
Hyperoxalate induces lipotoxicity and oxidative damage in NRK-52E cells. (A) Results of oil red staining observed by ordinary microscopy; red areas represent intracellular lipid accumulation. Oxalate concentrations: 0.5, 1, 1.5 mmol/L; treatment time: 24 h; (B) Expression of lipid metabolism-related proteins RXR, FABP4, PPARα, and oxidative damage protein Nrf2; (C) Semi-quantitative statistical plots of RXR, FABP4, PPARα, and Nrf2 protein expression; (D) semi-quantitative statistical plots of oil red staining area. Magnification is 200X, Scale bar = 100 μm; * indicates P < 0.05, *** indicates P < 0.001, ns indicates P > 0.05, no statistical difference. (E) Molecular models of HCA and Fenofibrate binding to protein PPARα, respectively; (F) Information on binding energy, number of H-bond binding receptors, and number of H-bonds for HCA and Fenobrate binding to PPARα, respectively.

### 3.6 Inhibition of oxalate-induced cellular lipotoxicity by HCA

To demonstrate the theoretical strength of the interaction between HCA and PPARα, we calculated the interaction force of HCA binding to PPARα using the Autodock docking tool, which can be used to simulate the docking of a drug to a target protein. We used fenofibrate, a specific agonist of PPARα, as a reference (binding energy of -4.59 kcal/mol, Figure 6E). Negative binding energy is generally considered to indicate that two molecules can spontaneously bond and interact. HCA forms 4 pairs of hydrogen bonds with residues of amino acid LYS-449 and 1 pair of hydrogen bonds with amino acid ARG-465 docked by PPARa, the binding energy of HCA to PPARa is -2.38 kcal/mol (Figure 6F). The binding energy of HCA to PPARα showed that HCA can spontaneously bind to PPARα with significant binding power, indicating that HCA can affect lipid metabolism-related pathways by binding to PPARα.

We then examined the effect of HCA on oxalate-induced lipotoxicity-related protein levels (Figure 7B). Oxalate treatment reduced the expression levels of PPARα, FABP4, and RXR proteins, while inhibiting the expression of antioxidant-related proteins HO-1, Nrf2, and SOD (p<0.05). The expression of PPARα, FABP4, and RXR was significantly restored under HCA treatment, while the intracellular antioxidant pathway was obviously activated. The accumulation of triglycerides is an important cause of lipotoxicity [22]. The triglyceride results showed that oxalate caused intracellular triglyceride accumulation (p<0.0001), and after HCA intervention, triglyceride levels in NRK-52E cells decreased by approximately 77.8% compared to the modeling group (p<0.0001, Figure 7E). The results of oil red staining also showed that HCA significantly reduced the accumulation of cellular lipids induced by high oxalate (Figure 7D).

**Figure 7.**
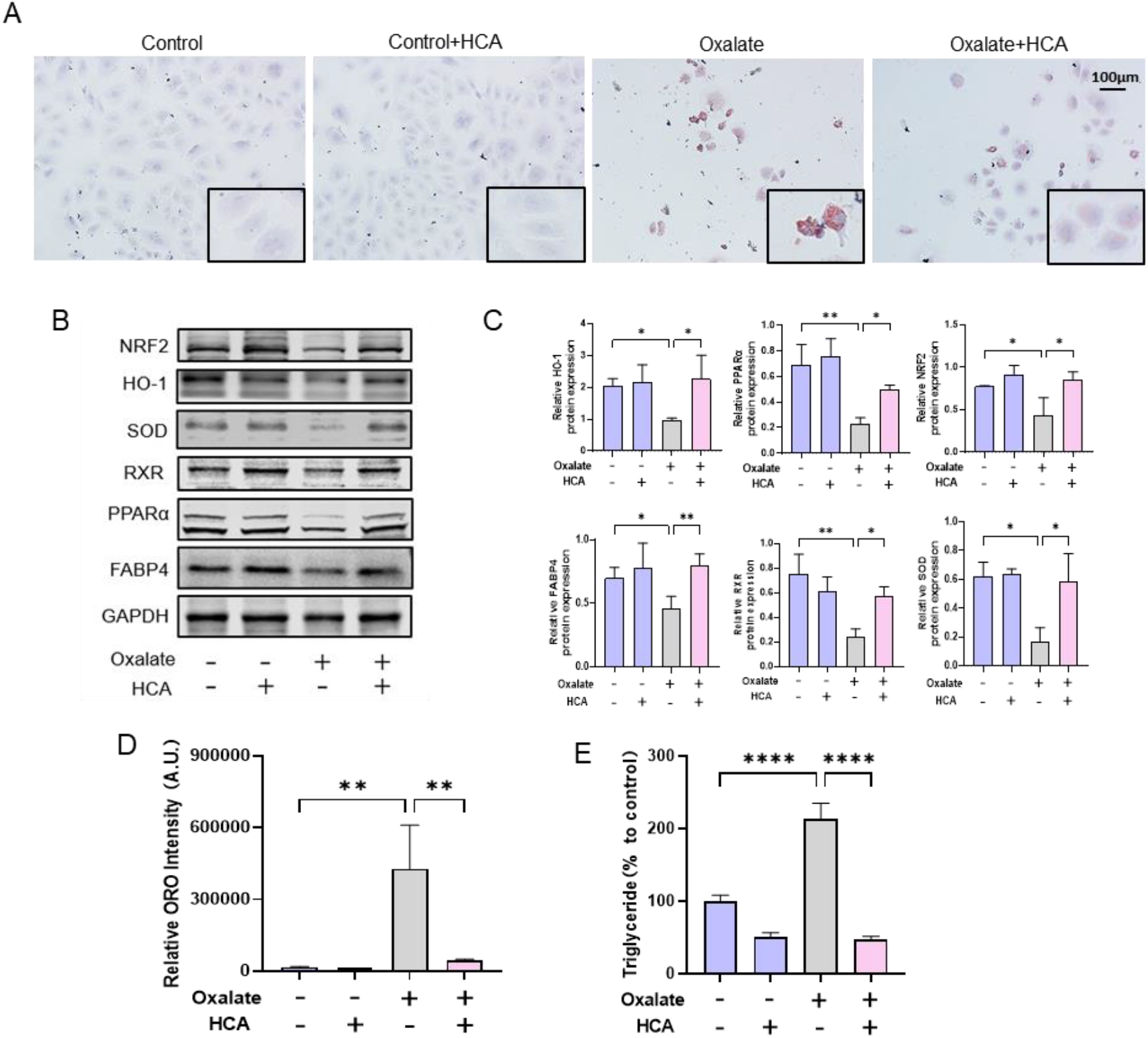
HCA inhibits cytosolic lipotoxicity induced by oxalate. (A) Microscopic observation of oil red staining, the red area represents intracellular lipid accumulation, magnification 200X, Scale bar=100 μm; (B, C) Expression of FABP4, PPARα, RXR, HO-1, Nrf2 and SOD proteins, and semi-quantitative statistical graphs; (D) Quantitative graph of intracellular triglycerides; (E) Semi-quantitative statistical graph of oil red staining. Oxalate concentration: 1.0 mmol/L; HCA concentration: 2.0 mmol/L; treatment time: 24 h. * indicates P<0.05, ** indicates P<0.01, and **** indicates P<0.0001.

We used PPARα-specific agonist Fenofibrate to activate PPARα and silence PPARα by associated siRNA, and examined PPARα pathway downstream related molecules and oxidative damage proteins (Figures 8E-8H). Fenofibrate significantly upregulated PPARα expression (p<0.001), and reduced oxalate-induced lipid toxicity (Figure 8A), which in turn elevated the expression of antioxidant molecules HO-1, Nrf2, and SOD (Figure 8E). In contrast, the activity of PPARα was inhibited by silencing treatment by associated siRNA, and the expression level was significantly decreased (p<0.001). Under this condition, HCA was unable to effectively upregulate the expression of PPARα and antioxidant molecules of Nrf2, HO-1, and SOD (Figure 8G), and was unable to reduce cellular lipotoxicity due to PPARa silencing (Figure 8C). PPARα is a key factor for HCA to inhibit the cytotoxicity caused by high oxalate. HCA could significantly reduce the lipotoxic damage caused by high oxalate to renal epithelial cells by activating the PPARα pathway.

**Figure 8.**
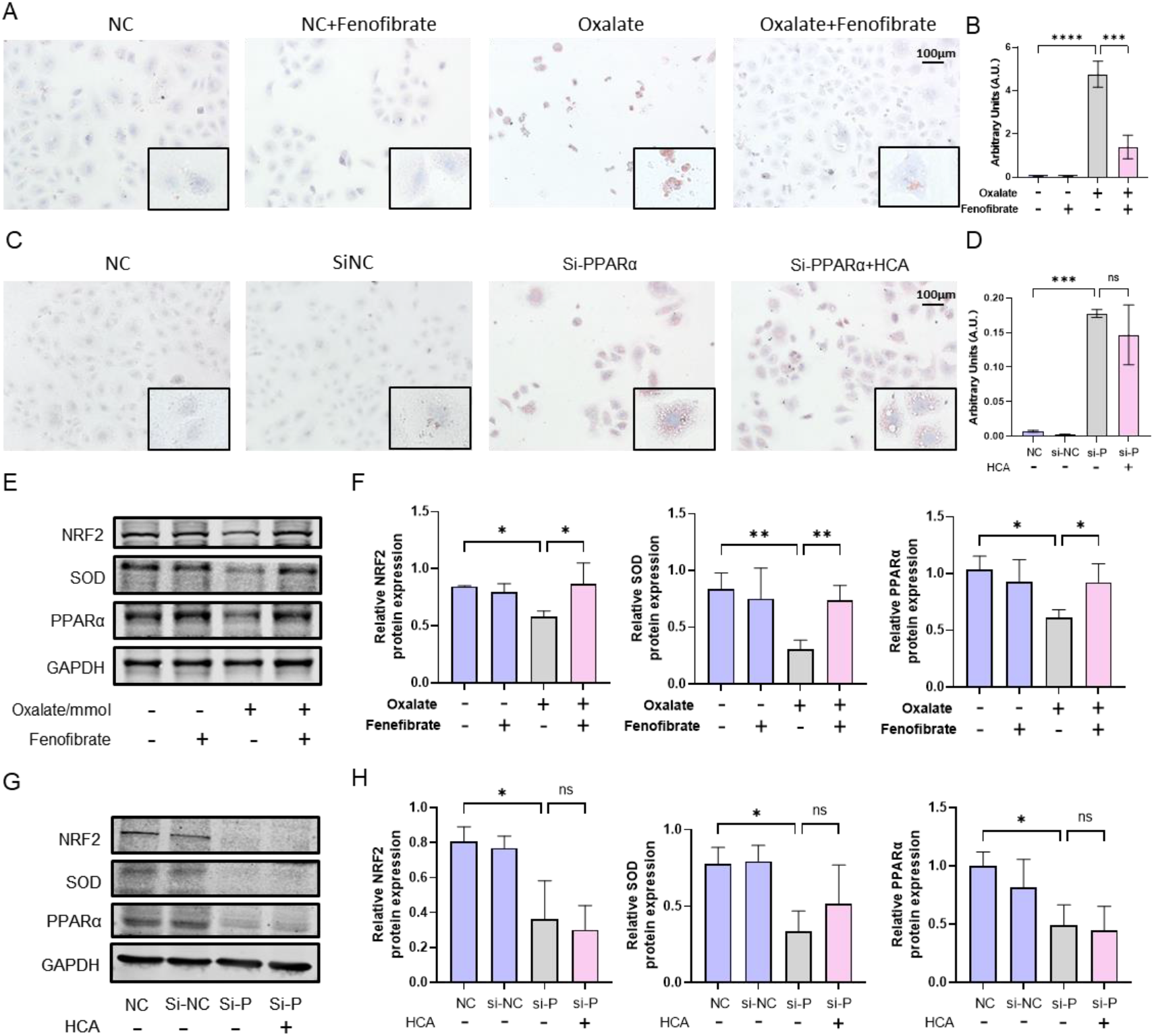
Effects of PPAR agonists and siPPARα on oxidative damage-related protein expression. (A&C) Microscopic observation of oil red staining, red area represents intracellular lipid accumulation, magnification 200X, Scale bar=100 μm; (E-H) PPARα, Nrf2 and SOD protein expression and semi-quantitative statistical graphs; (B&D) Semi-quantitative statistical graphs of oil red staining. Oxalate concentration: 1.0 mmol/L; Fenofibrate concentration: 1umol/L; HCA concentration: 2.0 mmol/L; Treatment time: 24 h. * indicates P<0.05, ** indicates P<0.01, *** indicates P<0.001, **** indicates P<0.0001.

## 4. Discussion

Hydroxycitric acid, a derivative of citric acid, is chemically similar to potassium citrate, a clinically used CaOx crystallization-inhibiting drug, and can inhibit the crystallization process of CaOx by competitively binding to Ca^2+^. Previous study reported that the binding energies of HCA, CA, and Ox complexing with Ca^2+^ ions is −174 kcal/mol, −144 kcal/mol, and −114 kcal/mol, respectively. The corresponding number of Ca^2+^ ions complexed per organic anion is 1.5, 1.5, and 1.0 [14]. Due to the increased hydrogen bonding in HCA, it shows a higher affinity for Ca^2+^ ion complexation relative to CA and Ox. HCA effectively inhibits the formation of up to 60% of COM crystals by reducing the rate of crystal growth and inhibiting COM crystal nucleation [14].

The primary urine formed by the filtration of blood takes only about 3-4 minutes to pass through the kidney unit [23]. Our results show that HCA significantly inhibits crystal growth within 30 min, with almost no significant CaOx crystal formation, and that the amount of crystallization does not increase very significantly with increasing crystallization time (Figure 3G). In addition, HCA also showed effective inhibition of CaOx crystallization in a high oxalate model in rats (Figure 1C). The stabilization and coating of early nanoparticles by HCA, together with the delay and reduction of bulk crystallization, should reduce crystallization and CaOx retention in the renal tubules. Furthermore, HCA can convert COM to COD crystals, which might be of therapeutic benefit due to the lower affinity of the latter to epithelial cell membrane molecules, lower cytotoxicity, and thus lower intratubular retention.

Interaction between cells and crystals, especially the adhesion of crystals to the cell surface, is the main cause of the rapid increase in stone size [24]. The increased oxalate concentration in urine leads to the migration of cell membrane components (e.g. phosphatidylserine) to the extracellular surface, causing high expression of negatively charged adhesion molecules such as osteopontin (OPN), hyaluronic acid, and CD44, which in turn promotes crystal-cell adhesion and exacerbates crystal-induced cell damage [25]. Cellular damage can promote CaOx adhesion on the cell surface, while reducing cellular damage by pharmacological intervention can significantly reduce crystal deposition and adhesion [26]. In the immunohistochemical results of rat kidneys, the expression levels of CD44 and OPN were significantly increased in the stone group, while the expression levels in the HCA group were similar to those of the normal group (Figure 1H). It is shown that HCA can obviously inhibit the expression of adhesion molecules and reduce the risk of crystal adhesion on the cell surface. In addition, the negatively charged HCA can adsorb onto the surface of CaOx crystals, increasing the electronegativity of the crystal surface and inhibiting the adhesion of CaOx crystals to the weakly negatively charged cell surface. In both animal and cellular experiments with HCA intervention, the number of crystals formed and adhered to the lumen and epithelial cell surface of the kidney tubules was significantly reduced (Figures 1C&3G) High concentrations of oxalate and excessive accumulation of CaOx crystals on the cell surface can cause cellular damage, resulting in cellular morphological disorders and skeletal rupture, causing an imbalance in mitochondrial membrane polarity and an increase in intracellular Ca^2+^ levels, leading to oxidative damage and ultimately cell death (Figures 4&5). Mitochondrial dysfunction may be an important event in oxalate-induced injury and plays a key role in the pathogenesis of kidney stones by activating intracellular ROS signaling pathways [27]. HCA may effectively inhibit the excessive production of intracellular ROS caused by oxalate, attenuate oxidative damage (Figures 5B&5C). In addition, HCA may complex the Ca^2+^ released from mitochondria, reducing intracellular Ca^2+^ damage to the cell (Figure 5A). Cell damage might promote the formation of CaOx crystals, leading to the crystallization of CaOx crystals at lower supersaturation levels, and greatly increases the adhesion and aggregation of formed crystals on the cell surface.

To reveal the molecular mechanism of HCA inhibition of oxalate nephropathy, RNA sequencing (RNA-seq) was performed on rat kidney tissue before and after HCA treatment (Figures 2A-2E). RNAseq results showed a similar gene expression profile between normal control and renal tissue treated with oxalate premixed with HCA, while oxalate alone induced drastic changes. Gene set enrichment analysis of differently expressed genes in the oxalate-treated group versus oxalate-treated group under HCA protection samples revealed oxalate-induced alterations of gene expression mainly involved in lipid metabolism-related pathways, inflammation, adhesion-related pathways, and oxidative stress-related pathways, with the most significant being the lipid metabolism-related PPAR pathway (Figure 2E). Therefore, we hypothesize that HCA may protect the kidney against oxalate-induced kidney injury by activating the PPAR signaling pathway. Studies show that patients with elevated dyslipidemia have an increased risk of developing kidney stone disease [28-30], Thus, lipids may play an important role in the development of kidney stone disease. Lipid content in the urine of patients with kidney stones has been reported to be positively correlated with the degree of tubular damage and oxidative stress [31]. Chao et al. [32] showed the presence of 196 different lipid metabolites in the kidney and serum of a mouse model of kidney stones, of which ceramide and lysophosphorylcholine mediated the inflammatory response and oil-based ethanolamine and glycerophosphoethanolamine acetal phospholipids induced oxidative stress, was closely associated with the development of kidney stones. Peroxisome proliferator-activated receptor alpha (PPARα) is a major regulator of lipid metabolism, involved in inflammation and oxidative stress, and can be activated by ω-3 fatty acids [33]. PPARα is a ligand-activated nuclear receptor and transcription factor that is a key regulator of fatty acid oxidation. PPARα promotes metabolic remodeling and lipid oxidation, and its dysregulated expression can lead to metabolic disorders in various organs of the body [34].

The accumulation of neutral lipids in non-adipose tissues can lead to various types of cell and organ damage, which in turn can lead to a variety of chronic diseases [10], including oxalate nephropathy[34-37]. The general pathogenic mechanism of lipotoxicity is thought to be an overload of intracellular free fatty acids, leading to an accumulation of intracellular triglycerides, which increases the production of ROS, apoptosis, or secretion or a combination of profibrogenic and proinflammatory factors [9, 10]. When lipid metabolism is blocked within the renal tubular cells, this can result in lipid accumulation leading to the development of lipotoxicity. The functional properties of the proximal tubule cell-dependent fatty acid oxidation pathway may be related to the distribution of PPARα in the kidney. PPARα is highly expressed in the renal proximal tubule, but rarely in the glomerulus and not at all in the loop of Henle, distal tubules, or collecting ducts [38].This distribution suggests that PPARα plays a key role in coordinating the energy metabolism of the renal tubular cells.

FABPs belong to a family of intracellular proteins that exhibit a high affinity for the non-covalent binding of long-chain fatty acids and transport endogenous fatty acids from the cell surface to various sites of intracellular fatty acid storage and metabolism[39]. PPARα has been reported to sense free fatty acids and their derivatives in the cytoplasm and subsequently transfer them to the nucleus. In the nucleus, PPARα forms a heterodimer with the Retinoid X Receptor (RXR), which can be activated by PPARα ligands alone or in concert, resulting in increased promoter activity of downstream related genes [34, 40]. Jao et al. [10] showed that downregulation of PPARa inhibited the mitochondrial fatty acid oxidation pathway, which in turn led to lipid accumulation in renal tubular epithelial cells, ultimately leading to tubular epithelial cell dysfunction. Chen et al. [41] showed that fenofibrate could reduce the expression of pro-inflammatory factors, inhibit the NF-κB pathway, significantly improve lipid distribution, and inhibit tubulointerstitial fibrosis and interstitial macrophage infiltration by activating PPARα.

Our results showed that oxalate stimulation significantly inhibited the PPARα pathway thereby causing oxidative damage to cells, causing a significant increase in ROS levels, triglyceride levels, and the number of oil-red positive stained cells (Figures 6A-6C). High oxalate stimulation attenuated cytoplasmic FABP4 and PPARα expression in NRK-52E, resulting in blockage of the intracellular fatty acid transport pathway, which led to the inability of mitochondria to use fatty acids normally for energy supply. At the same time, the expression of ACOX1 in mitochondria decreased, the fatty acid β-oxidation pathway was impaired, and the expression of antioxidant proteins Nrf2, SOD, and HO-1 decreased, activating the ROS pathway and ultimately causing oxidative damage to the cells. At the same time, HCA can also protect the kidney by activating the PPARα pathway to reduce oxalate-induced lipotoxicity levels and activating the expression of antioxidant molecules such as Nrf2 and SOD (Figure 9).

**Figure 9.**
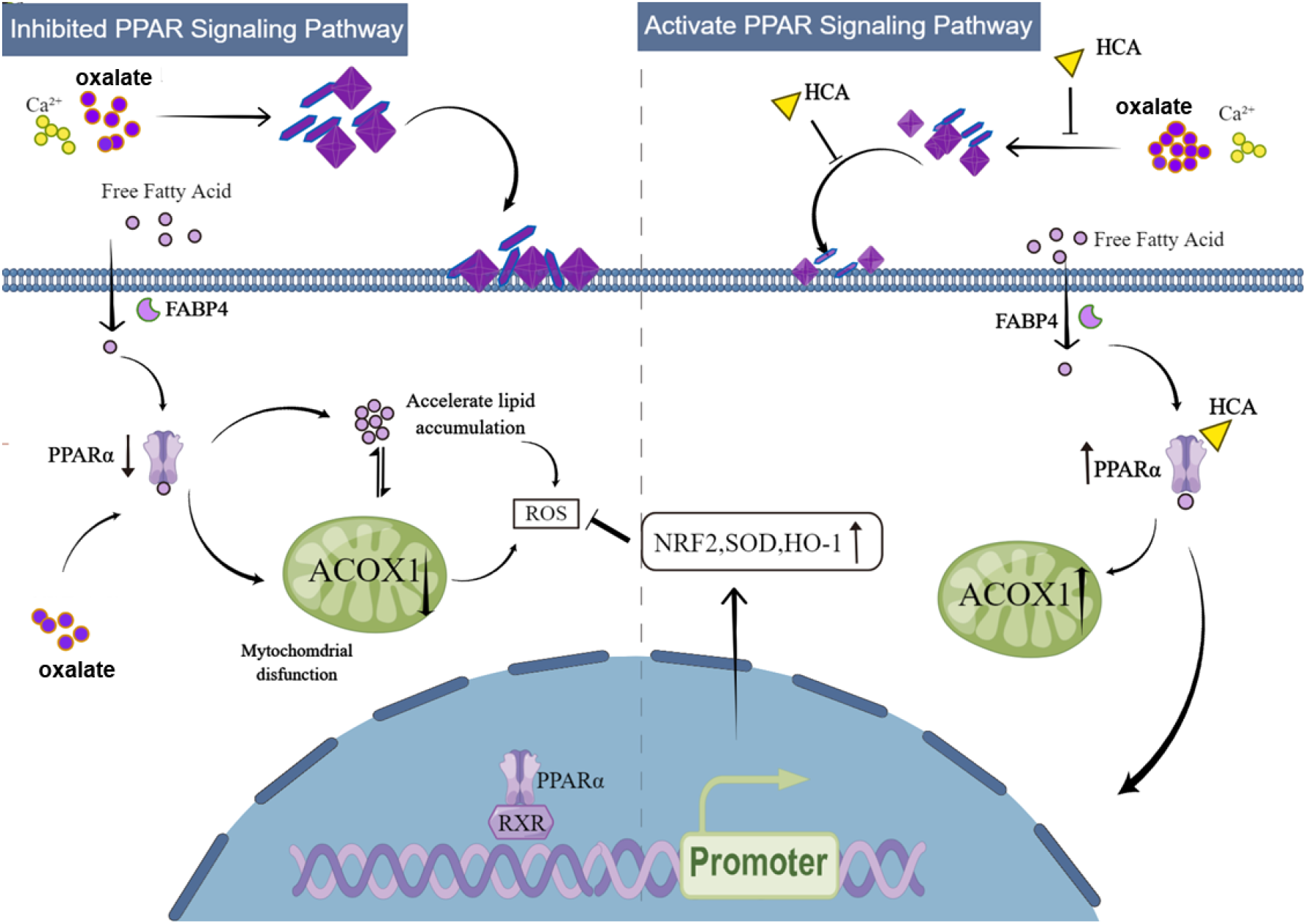
HCA attenuates lipid accumulation and subsequent oxalic nephropathy by inhibiting CaOx formation and regulating the PPARα pathway

Fenofibrate as a PPARα agonist also interfered with oxalate damage-induced lipotoxicity levels (Figure 8A), further validating that HCA plays a similar role to fenofibrate in activating PPARα to reduce lipotoxicity. We further used siPPARα to transfect NRK-52E cells and found that reducing PPARα protein expression could increase the level of lipotoxicity (Figure 8C), which in turn led to a decrease in the expression of oxidative stress-related molecules Nrf2, SOD, and HO-1 (Figure 8G), causing oxidative damage. The above results suggest that HCA can enhance intracellular antioxidant levels through activation of the PPARα pathway, thereby resisting oxidative damage caused by high oxalate.

In conclusion, we found that HCA is a novel drug that can inhibit oxalate-induced lipotoxicity and CaOx crystal formation, and therefore may provide new opportunities for the pharmacological treatment of oxalate nephropathies.

## Acknowledgements

This work was granted by the Science and Technology Plan Project of Guangzhou (NO. 202102010306, 202102010165), National Natural Science Foundation of China (NO. 21975105), and Guangdong Provincial Science and Technology Plan Project (No. 2017B030314108).

